# Antagonistic antimalarial properties of a methoxyamino chalcone derivative and 3-hydroxypyridinones in combination with dihydroartemisinin against *Plasmodium falciparum*

**DOI:** 10.1101/2022.11.19.517177

**Authors:** Tanyaluck Kampoun, Pimpisid Koonyosying, Jetsada Ruangsuriya, Parichat Prommana, Philip J. Shaw, Sumalee Kamchonwongpaisan, Hery Suwito, Ni Nyoman Tri Puspaningsih, Chairat Uthaipibull, Somdet Srichairatanakool

**Affiliations:** Department of Biochemistry, Faculty of Medicine, Chiang Mai University, Chiang Mai, Thailand; Medical Molecular Biotechnology Research Group, National Center for Genetic Engineering and Biotechnology (BIOTEC), Thailand Science Park, Pathum Thani, Thailand; Department of Chemistry, Faculty of Science and Technology, Airlangga University, Surabaya, Indonesia; Laboratory of Proteomics, University-CoE Research Center for Bio-Molecule Engineering, Universitas Airlangga, Kampus C-UNAIR, Surabaya, East Java, Indonesia

**Keywords:** *Plasmodium*, antimalarial, artemisinin, chalcone, hydroxypyridinone, drug resistance, ferredoxin

## Abstract

The spread of artemisinin (ART)-resistant *Plasmodium falciparum* threatens the control of malaria and mutations in the propeller domains of *P. falciparum* Kelch13 (*k13*) are strongly associated with the resistance. Ferredoxin (Fd) in the ferredoxin/NADP^+^ reductase (Fd/FNR) redox system is essential for isoprenoid precursor synthesis in the plasmodial apicoplast; nonetheless, mutations of Fd gene (*fd*) may modulate ART resistance and Fd would be an important target for antimalarial drugs. We investigated the inhibitory effects of dihydroartemisinin (DHA), methoxyamino chalcone (C3), and iron chelators including deferiprone (DFP), 1-(*N*-acety1-6-aminohexyl)-3-hydroxy-2-methylpyridin-4-one (CM1) and deferiprone-resveratrol hybrid (DFP-RVT) against the growth of wild-type (WT) *P. falciparum* parasites and those with *k13* and *fd* mutations. C3 showed antimalarial potency similar to the iron chelators. Surprisingly, combined treatments of DHA with the C3 or iron chelators showed moderately antagonistic effects against P. falciparum growth. No differences were observed among the mutant parasites with respect to their sensitivity to C3 and the chelators, or the interactions of these compounds with DHA. The data suggest that inhibitors of the Fd/FNR redox system should be avoided as ART partner drugs in ART combination therapy for treating malaria.

## Introduction

Artemisinin (ART)-based combination therapy is recommended by the World Health Organization for the first-line treatment of malaria in uncomplicated *Plasmodium falciparum* infections. ART and its derivatives are bio-transformed by the liver to the active metabolite, dihydroartemisinin (DHA) (Figure 1), which can be activated by iron, resulting in endoperoxide radicals that damage proteins via the formation of covalent adducts. The accumulation of damaged, polyubiquitinated proteins rapidly induces lethal endoplasmic reticulum stress [1]. ART-resistant *P. falciparum* parasites emerged in Western Cambodia in the 2000s showing a slow clearance phenotype in malaria patients and an increased ring survival rate [2,3]. Mutations in the kelch domain of the Kelch13 (*k13*) gene have been established as genetic markers for ART-resistant malaria parasites [4,5]; however, ART resistance is due to other genetic factors in some resistant parasites that do not contain *k13* mutations [6,7]. A genome-wide association study identified variants in the ferredoxin *fd*), apicoplast ribosomal protein S10 (*arps10*), multidrug resistance protein 2 (*mdr2*), and chloroquine resistance transporter (*crt*) genes as additional factors contributing to ART resistance, in which a missense *fd* mutation *fd*-D193Y) was the most frequent variant among resistant parasites [8].

**Figure 1:**
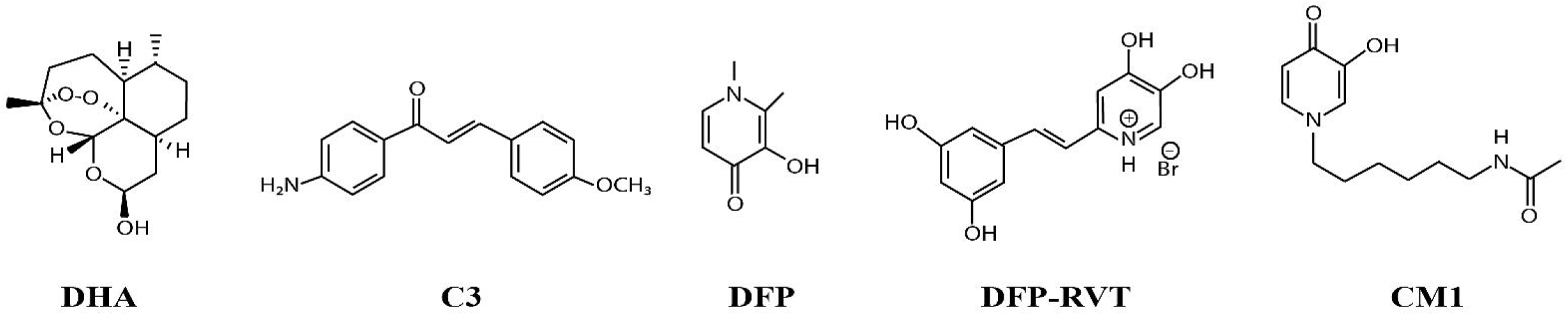
Chemical structures of dihydroartemisinin (DHA), compound 3 (C3), and 3-hydroxypyridinones; deferiprone (DFP), deferiprone-resveratrol (DFP-RVT), and 1-(*N*-acetyl-6-aminohexyl)-3-hydroxy-2-methylpyridin-4-one (CM1).

Ferredoxin (Fd) is an iron-sulfur (Fe-S) electron carrier protein that functions with ferredoxin NADP^+^ reductase (FNR) in apicoplast metabolism, particularly in isoprenoid biosynthesis, fatty acid desaturation, and heme oxygenation. However, only the isoprenoid biosynthesis pathway is essential during intraerythrocytic developmental stages [9,10]. The Fd-D97Y mutant protein (mutated residue corresponding to residue 193 in the full-length preprocessed Fd protein) has increased binding affinity for FNR. This mutation suppresses FNR function; hence, loss-of-function mutations in the Fd/FNR system could contribute to ART resistance [11]. However, Fd/FNR mutations are constrained because this redox system is essential and a validated antimalarial drug target [12,13]. The Fd-FNR interaction is inhibited in vitro by methoxyamino chalcone derivatives, of which the most potent compound, C3 (Figure 1), exhibits 50% inhibition at 100 μM [14].

In addition to Fd, iron is a cofactor for enzymes such as aconitase, oxidoreductases, and ribonucleotide reductase. Moreover, hemoglobin, myoglobin, and cytochromes contain an iron-prosthetic heme group. Iron also acts as a chemical catalyst for the generation of reactive oxygen species (ROS) via the Fenton and Haber-Weiss reactions. Iron chelators, such as desferrioxamine (DFO), 1,2-dimethyl-3-hydroxypyridin-4-one or deferiprone (DFP) and deferasirox (DFX) are clinically used for the treatment of patients with iron overload. Importantly, DFO, DFP, DFX, and other iron chelators, including 1-(*N*-acety1-6-aminohexyl)-3-hydroxy-2-methylpyridin-4-one (CM1) and deferiprone-resveratrol hybrid (DFP-RVT) possess antimalarial activity by interfering with iron uptake, depleting the intraerythrocytic labile iron pool in parasites and expediting the phagocytosis of ingested parasites [15-20]. Some antimalarial drugs, such as quinine, mefloquine, and artesunate are antagonized by DFP [21].

In combination therapy, two different drugs are used to increase the efficacy of treatment, and the interaction of these drugs can be assessed by isobologram analysis [22]. Drugs with a connection in their mechanisms of action, for example, those acting on the same metabolic pathway, can show synergistic or antagonistic interactions. For example, iron chelators are antagonistic to ART and related compounds [14,23], which is explained by the iron-dependent activation of ART [21,24,25]. Although the putative loss of function in the *fd*-D193Y mutant is associated with ART resistance, it does not affect the survival of *P. falciparum* parasites exposed to DHA [26,27]. Therefore, it is unclear whether the parasite genetic background, in particular, the allelic status of the *fd* and *k13* genes affects the antimalarial potency of compounds targeting Fd/FNR and/or the interaction of these compounds with DHA. Currently, we have identified growth inhibition in the ART-sensitive *P. falciparum* 3D7 strain and genome-edited lines of this parental strain with fd and k13 mutations (fd^D193Y^_3D7, k13^C580Y^_3D7, and k13 ^C580Y^fd^D193Y^_3D7 [27]. We hypothesized that the antimalarial potency of chalcone derivatives C3 and our iron chelators and their interactions with DHA would be affected by the genetic background of the parasite. Thus, we focused our attentions on evaluating the potencies of C3, DFP, DFP-RVT and CM1 monotherapy and the combinations with DHA against the growth of wild-type (WT) *P. falciparum* parasites and those with *k13* and *fd* mutations.

## Materials & Methods

### Chemicals and Reagents

D-Sorbitol, dimethylsulfoxide (DMSO), hydroxyethylpiperazine ethane sulfonic acid (HEPES), RPMI 1640 medium and SYBR^®^ Green I were purchased from Sigma-Aldrich Chemical Company (St. Louis, MO, USA). Milli^®^-deionized water (DI) was purchased from Merck KGaA (Darmstadt, Germany).

### Drug and Compounds

DHA (item No. 19846, MW = 284.4 g/mol) was purchased from Cayman Chemical Company (Ann Arbor, MI, USA). C3 compound (MW = 254.1183 g/mol) was synthesized by and kindly supplied by Dr. Hery Suwito [14]. DFP (MW = 139 g/mol) was purchased from Sigma-Aldrich (St Louis, MO, USA). DFP-RVT (MW = 340 g/mol) was kindly supplied by Dr. Yongmin Ma, School of Pharmaceutical and Chemical Engineering, Taizhou University, Taizhou, People’s Republic of China, which was synthesized by Xu and colleagues [28]. 1-(N-Acetyl-6-aminohexyl)-3-hydroxy-2-methylpyridin-4-one or CM1 (MW = 266 g/mol) which is a DFP analogue and an orally active bidentate iron chelator, was synthesized and kindly supplied by Dr. Kanjana Pangjit, College of Medicine and Public Health, Ubon Ratchathani University, Ubon Ratchathani, Thailand [17,29].

### Stock Solutions of Compounds

For the parasite growth inhibition assay, stock solutions of DHA, C3 compound, and DFP-RVT were prepared with 1% (*w/v*) DMSO as the solvent, whereas DFP and CM1 were dissolved in sterile DI water at 1,000 times the highest dose tested. Ten two-fold serial dilutions were prepared from the stock solutions. For the drug combination study, the drug or compounds were first dissolved in solvents at 2,000 times the highest dose tested. Five two-fold serial dilutions of the stock solutions were prepared. Stock solutions were sterilized using a sterile syringe filter (hydrophilic polyvinylidene difluoride membrane, 0.22 μm pore size, Sigma-Aldrich Chemicals Company, St. Louis, MO, USA) and stored at −20 □C.

### *Plasmodium falciparum* Culture and Synchronization

The 3D7 parasite line and the transgenic 3D7 parasite lines in this study, which were established in our earlier study, were cultured and maintained in the same conditions in which the parasites grew at the same rate in these culture conditions [27]. Briefly, the parasites were cultured in complete RPMI 1640 medium pH7.4 containing 2 mM L-glutamine, 25 mM HEPES, 2 g/L NaHCO_3_, 27.2 mg/L hypoxanthine, and 0.5% Albumax II using human O^+^ blood group erythrocytes (2-4% hematocrit). Parasite cultures were incubated in 90% N_2_, 5% CO_2_, and 5% O_2_ at 37 □C and synchronized to the ring stage using 5% D-sorbitol treatment. Parasite developmental stage and viability were routinely assessed by microscopic examination of Giemsa-stained thin smear films.

### *Plasmodium falciparum* Growth Inhibition Assays

Before use, stock solutions of the compounds were 100X diluted in a complete RPMI 1640 medium. Ten microliters of each dilution were added to each well of a black 96-well plate. The synchronized ring-stage *P. falciparum* culture was suspended in complete RPMI 1640 medium to achieve the parasite suspension with 1% parasitemia and 2% hematocrit, which was dispensed 90 μL/well to perform the treatments. The parasite was treated in technical duplicates with various doses of the test compound in a black 96-well plate, while a mock treatment (parasite suspension in culture medium with 0.1% DMSO) was also performed. The mock treatment was assigned as 100% parasite growth. The treated parasites were incubated under standard culture conditions for 48 h and the parasite survival rate was determined using the malaria SYBR Green I-based fluorescence (MSF) assay as described previously [16].

For the MSF assay, 0.2 μL of SYBR^®^ Green I was diluted in 1 mL of lysis buffer solution containing 20 mM Tris-HCl, 5 mM EDTA, 0.008% (*w/v*) saponin and 0.08% (*v/v*) Triton X100. Then, 100 μL of SYBR^®^ Green I solution was added to each well of a 96-well black microplate (Corning^®^ Product number CLS 3601, polystyrene flat-bottom type, Merck KGaA, Darmstadt, Germany). The plate was mixed using a MixMate^®^ Eppendorf machine (Eppendorf SE, Hamburg, Germany) at 1,000rpm for 30 s and incubated for 1 h in the dark at room temperature. Fluorescence intensity (FI) was measured using a fluorescence multi-well plate reader (Beckman Coulter AD340C, Beckman Coulter Inc., Brea, CA, USA) with excitation and emission wavelengths of 485 and 530 nm, respectively. The concentration of the test compound with 50% inhibition of growth (IC_50_) was calculated using R program with drc R package [30].

### Combination Treatment against *P. falciparum* Growth

Working solutions of the drug or compound were mixed in a 1:1 ratio (5 μL each) in a checkerboard manner in a black 96-well plate, in which the concentration of one drug or compound was fixed while the concentrations of the other were varied. The parasite suspension (1% rings, 2% hematocrit) was then added to each well (100 μL/well in total) and cultured for 48 h under standard conditions. The parasite growth was determined using the MSF assay as described above. Data were obtained from at least three independent replicates, with two technical replicates per experiment.

### Isobologram Analysis of Drug Combination

The interaction of DHA with Fd-FNR inhibitor or iron chelator was assessed by isobologram analysis of IC_50_ values. The FIC indices were calculated from the ratio of the IC_50_ value obtained from the combination treatment to that obtained from the single compound treatment. The □FIC for the combination is the summation of an individual drug or compound. The FIC value was calculated as follows:

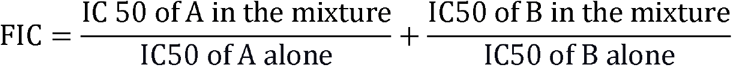

The FICs of the combinations from at least three individual experiments were used to calculate the mean values of □FIC. Interactions were assigned as synergistic (□FIC <0.9), additive (0.9 < □FIC < 1.1), or antagonistic (□FIC >1.1), with 1.1 < □FIC < 1.2 representing slight antagonism, 1.2 < □FIC < 1.45 representing moderate antagonism, 1.45 < □FIC < 3.3 representing antagonism, 3.3 < □FIC < 10 representing strong antagonism, and □FIC >10 representing very strong antagonism [31,32].

### Statistical Analysis

Data for each experiment were obtained from at least three independent replicates. The IC_50_ values of the mutant parasites were compared with those of the parental 3D7 parasite using drc R package by pair-wise t-tests with Bonferroni-Holm post-hoc correction of the p-values, in which p <0.05 was considered significant [30]. The mean □FICs of *fd*-D193Y parasites were statistically compared with the mean □FIC of fd WT parasites using GraphPad Prism 8.3.0 software by unpaired t-test with Welch’s correction.

## Results

### Single Compound Sensitivity Test of *P. falciparum*

The growth inhibitory properties of DHA, C3, DFP, DFP-RVT and CM1 were assessed in *P. falciparum* parental 3D7 and transgenic parasites fd^D193Y^_3D7, k13^C580Y^_3D7, and k13C^580Y^fd^D193Y^_3D7. The potency of the compounds varied as follows: DHA > DFP-RVT > C3 > DFP ≈ CM1. However, the IC_50_ values of the compounds were not significantly different between the parental and transgenic strains (Figure 2).

**Figure 2:**
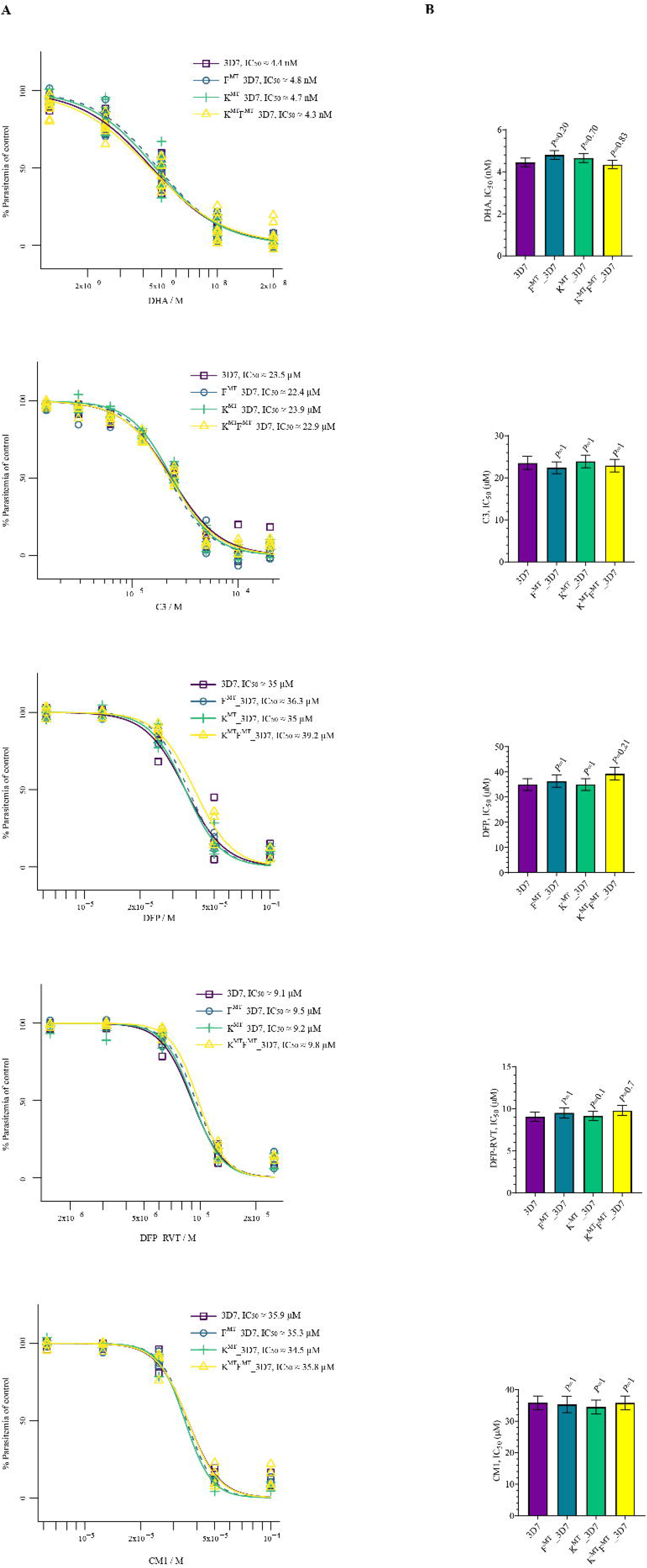
Dose-response analysis of growth inhibition for DHA, C3, DFP, DFP-RVT, and CM1. *P. falciparum* parental parasite 3D7 and transgenic parasites fd^D193Y^_3D7, k13^C580Y^_3D7, and k13^C580Y^ fd^D193Y^_3D7 were treated with varying doses of each compound. (A) Data were obtained from at least three independent experiments and dose-response models were fitted to the data. (B) The IC_50_ values and the associated confidence intervals are shown. *P*-values from pair-wise t-test with Bonferroni-Holm post-hoc correction comparing the IC_50_ of each transgenic parasite to that of the parental 3D7 parasite are shown above the IC_50_ values. F^MT^ = fd^D193Y^ mutation, K^MT^ = k13^C580Y^ mutation.

### Test of Pharmacological Interaction between DHA and Test Compounds in *P. falciparum*

The interactions between DHA and the test compounds were assessed as the fractional inhibition concentration (□FIC) index, and the mean □FICs of the combinations were compared between *fd* wild-type (fd^WT^) and *fd*-mutated (fd^MT^) parasites using Welch’s T-test (Figure 3). An antagonistic effect was indicated when □FIC >1.1, an additive effect was indicated when 0.9 < □FIC < 1.1, and synergism was indicated when □FIC< 0.9, using the criteria established by previous studies of antimalarial interactions [31,32]. For the DHA and C3 combinations, the mean □FICs were moderately antagonistic. Moreover, no significant differences in mean □FIC were observed different between fd^WT^ and fd^MT^ parasites. All combinations of DHA with iron chelator compounds were antagonistic, although no significant difference was observed between fd^WT^ and fd^MT^ parasites.

**Figure 3.**
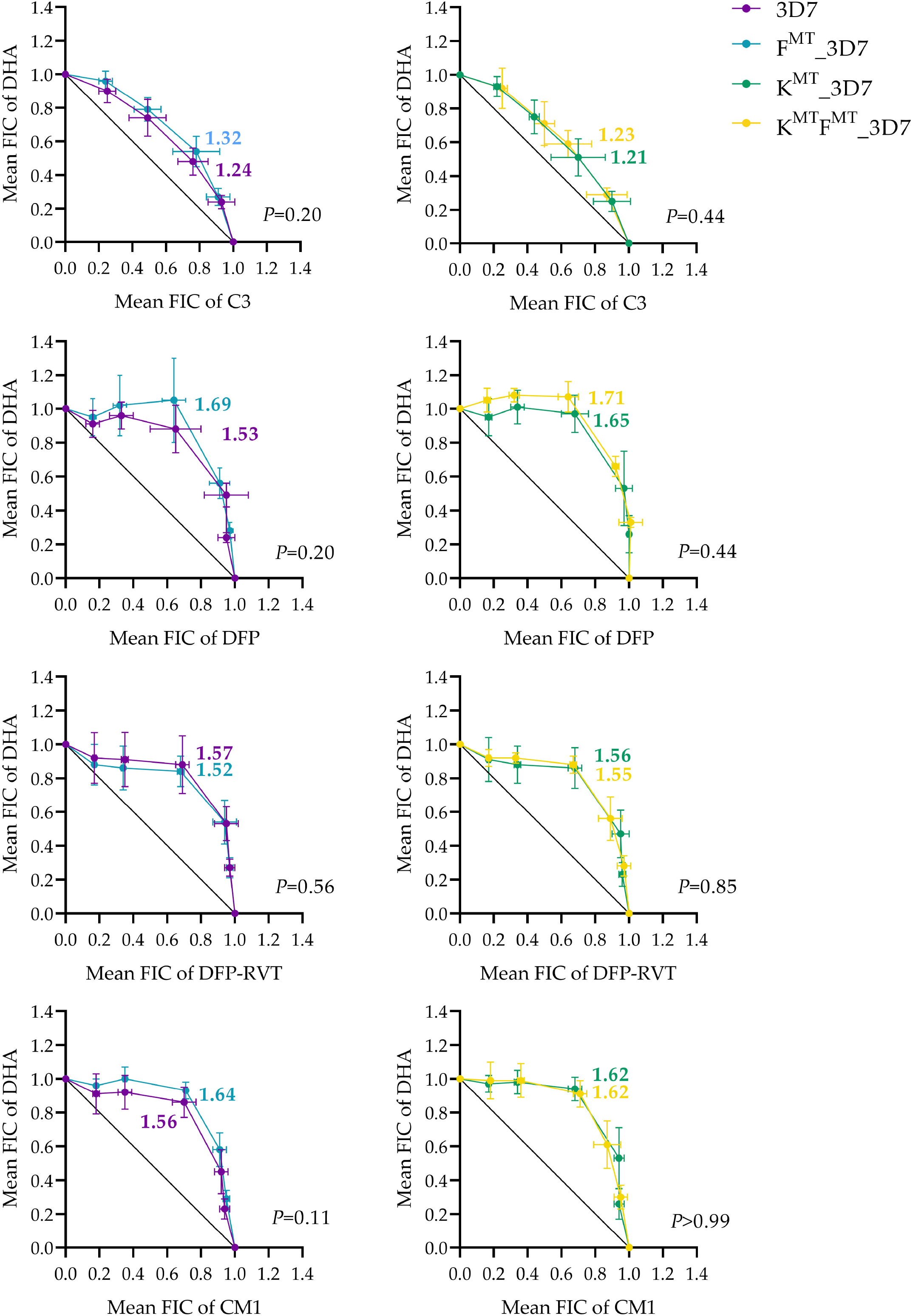
Isobolograms for the interaction between DHA and test compounds (candidate Fd/FNR inhibitor C3 or iron chelators DFP, DFP-RVT, and CM1) against the parental 3D7 and transgenic F^MT^_3D7 (*fd* mutant, *k13* wild-type), K^MT^_3D7 (*fd* wild-type, *k13* mutant), and K^MT^F^MT^_3D7 (*fd* mutant, *k13* mutant) *P. falciparum*. Data were obtained from at least three independent experiments; the values shown are mean and the error bars represent SD. The ΣFIC values for each combination are shown next to the isobologram. *P*-values for the *t*-test of the pairwise comparison of ΣFIC values between *fd*^MI^ and *fd* wild-type parasites are shown on each graph. F^MT^ = fd^D193Y^ mutation, K^MT^ = k13^C580Y^ mutation.

## Discussion

Our finding that C3 antagonizes ART is in line with a recent report that other apicoplast-targeting antimalarials also antagonize ART [31]. These findings caution against the use of ART partner drugs that inhibit the Fd-FNR interaction and other apicoplast targets, which may reduce ART efficacy in malaria chemotherapy. The antimalarial potency of the Fd-FNR inhibitor C3 was modest (IC_50_ range 23-25 μM) and did not differ between 3D7 and genome-edited parasites, including those with the Fd D193Y mutation. From the Fd docking study, C3 was predicted to bind Fd at a site distant from the D193Y mutation [14], which might explain why no difference in antimalarial potency was observed between fd^WT^ and fd^MT^ parasites. In contrast, the antimalarial effect of C3 could be due to the inhibition of targets other than Fd, since the antimalarial IC_50_ is markedly lower than the concentration of C3 required for 50% inhibition of the interaction between recombinant Fd and FNR proteins (100 μM) [14]. The potential for promiscuous targeting by the chalcone derivative C3 is plausible since other chalcone derivatives such as licochalcone A are known to be promiscuous-targeting antimalarials. Licochalcone A inhibits *P. falciparum* mitochondrial complexes II and III, and possibly the erythrocyte membrane [33,34]. In this study, the tested iron chelators showed modest antimalarial activity, similar to previous reports [15,16,18,35]. No differences in sensitivity to DHA were observed among the parasites tested, although it should be noted that the ART resistance phenotype manifested in *k13* mutants is not detectable by the growth inhibition assay employed in this study [5].

Importantly, the interactions of C3 with DHA in different parasite backgrounds were moderately antagonistic; hence, the inhibition of the Fd-FNR interaction by C3 could antagonize DHA. Nevertheless, there was no significant difference between the fd^WT^ and fd^MT^ parasites and the antimalarial mode of action of C3 remain inconclusive at this point. Expectedly, the iron chelators were found to antagonize the antimalarial activity by DHA. Since no significant differences in the mean ΣFICs between fd^WT^ and fd^MT^ parasites were observed, it can be concluded that this *fd* mutation may not affect the antagonistic interaction between DHA and iron chelators. Presumably, *fd* mutations associated with the loss of Fd function have negligible impact on the iron pool responsive to iron chelators. The effect of the *fd* mutation on the interaction with ART might be smaller than that of C3, such that it cannot be detected by our experimental approach.

## Conclusions

The putative Fd-FNR inhibitor C3 demonstrated antimalarial activity comparable to that of the iron chelators DFP, DFP-RVT, and CM1. No difference in sensitivity was observed among parasites with mutations in the *fd* and *k13* genes compared with the parental *P. falciparum 3D7* strain for any compound tested. In combination with DHA, C3 showed moderate antagonistic interactions, and the iron chelators were antagonistic. Other approaches and additional experimentations are required to understand the effect of the *fd* mutation on the Fd function and how this affects the interaction with ART.

## Acknowledgements

We express our sincere thanks to Dr. Yongmin Ma and Dr. Kanjana Pangjit for supplying DFP-RVT and CM1 compounds, respectively. We also thank the blood donors who supplied blood for *P. falciparum* culture.

## Additional Information and Declarations

## Abbreviations

ANOVA: analysis of variance
*arps10*: apicoplast ribosomal protein S10 gene
ART: artemisinin
CI: confidential interval
CM1: 1-(*A*-acetyl-6-aminohexyl)-3-hydroxy-2-methylpyridin-4-one
*crt*: chloroquine-resistant transporter gene
C3: Fd-FNR inhibitor compound 3
DFO: desferrioxiamine
DFP: deferiprone
DFP-RVT: deferiprone-resveratrol hybrid
DFX: deferasorix
DHA: dihydroartemisinin
DI: deionized water
DMSO: dimethyl sulfoxide
Fd: ferredoxin
*fd*: ferredoxin gene
*fd*-D193Y: *fd* mutation
Fe-S: iron-sulfur
FI: fluorescence intensity
FIC: fractional inhibition concentration
FIC: fractional inhibition concentration index
FNR: ferredoxin NADP^+^ reductase
HEPES: hydroxyethylpiperazine ethanosulfonic acid
IC_50_: inhibitory concentration 50%
K13: Kelch 13
*k13*: Kelch 13 gene
*k13*^C580Y^ mutant and *k13*^MT^fd^MT^ mutant: k13 gene mutant
*mdr2*: multidrug resistance protein 2
*P. falciparum*: *Plasmodium falciparum*
MSF: malaria SYBR Green I-based fluorescence
ROS: reactive oxygen species.

## Author Contributions

Conceptualization, C.U., S.K. and S.S.; methodology, C.U., P.P., S.K. and T.K.; investigation, P.K. and T.K.; resources, H.S. and N.P.; writing—original draft preparation, T.K. and S.S.; writing—review and editing, S.S.; visualization, J.R. and P.K.; supervision, C.U., P.S. and S.S.; project administration, C.U. and S.S.; funding acquisition, C.U. and S.S. All authors have read and agreed to the published version of the manuscript.

## Funding

This research was funded by the Royal Golden Jubilee PhD. Program (NSTDA), Thailand Research Fund (PHD0234/2558) and the Faculty of Medicine Endowment Fund, Chiang Mai University (grant number 113-2561). The APC was funded by the Royal Golden Jubilee PhD. Program (NSTDA), Thailand Research Fund (PHD0234/2558).

## Human Ethics

The study protocol was approved by expedited review (Project Code NIRB-024-2561).

## Data Availability Statement

All data are presented in figures.

## Conflicts of Interest

The authors declare no conflict of interest. The funders had no role in the design of the study; in the collection, analyses, or interpretation of data; in the writing of the manuscript; or in the decision to publish the results.

## Notes

### Competing Interest Statement

The authors have declared no competing interest.

